# A prioritization strategy for protecting Conservation Imperatives Sites

**DOI:** 10.64898/2026.05.01.721008

**Authors:** Joe Gosling, Eric Dinerstein, Anup R. Joshi, Neil D. Burgess, Haley Mellin, Lucas Joppa, Karl Burkart, Heather C. Bingham, Osgur McDermott-Long, Jasmin Upton

**Affiliations:** UNEP-WCMC; Conservation X Labs; University of Minnesota; Conserve.org; Haveli Investments

**Keywords:** threatened species, irreplaceability, conversion pressure, protected areas, OECMs, Target 3, Target 4, KMGBF

## Abstract

To prevent species extinctions, targeted action must focus on areas of threatened biodiversity facing intense human pressures. This objective is even more important in the run-up to 2030, the target date to conserve 30% of lands and waters globally. Conservation Imperatives (unprotected terrestrial sites that harbour rare, range-restricted, and threatened species) are critical to preventing imminent species losses. To prioritize among the 16,825 Conservation Imperatives Sites spanning 1.64 million km^2^, we ranked each site using a prioritization framework based on four criteria: number of threatened species per site; irreplaceability of the site; the proportion of an ecoregion’s remaining habitat contained in the site; and conversion pressure. Our approach prioritizes 1,667 sites representing 501,426 km^2^, or 0.37% of Earth’s terrestrial surface, most in need of urgent protection, with 87.34% of these sites occurring in 20 countries and in 250 ecoregions. This prioritization directly addresses the concern that protected areas must be targeted to protect endangered species, habitats and populations: 33.46% of the prioritized Conservation Imperatives Sites scored higher in irreplaceability than 90% of existing protected areas. Additionally, 51.53% are within 2.5 km^2^ of an existing protected area, making extending protection or restoring connectivity more feasible. Targeting conservation actions, especially in this small set of countries and ecoregions identified here, would contribute “high quality” areas for biodiversity as part of reaching the 30% coverage target by 2030.

## Introduction

Biodiversity is under immense human pressure and is declining faster now than at any point in human history (IPBES, 2019). Approximately one million species are projected to face extinction (IPBES, 2019; Tollefson, 2019), a figure that may itself be an underestimation (Liu et al., 2022). Compounding this crisis, human-induced pressures continue to intensify: almost 23% of land may face conversion by 2030, potentially affecting 460 million ha of intact natural lands (Oakleaf et al., 2024).

Conservation Imperatives Sites (unprotected areas harbouring rare, range-restricted, and threatened species) have been proposed as priority areas for protection to prevent the most likely and imminent extinctions (Dinerstein et al., 2024). Given that resources, time, and capacity are limited, prioritizing protection efforts among the 16,825 Conservation Imperatives Sites identified by Dinerstein et al. (2024) is essential to coordinate responses from multilateral and bilateral donors, host-country governments and NGOs, foundations, the private sector, local community organisations and civil society to the sites requiring urgent attention. Here, we introduce a framework to prioritize the existing set of Conservation Imperatives Sites based upon four criteria: 1) the number of threatened species per site; 2) irreplaceability of the site; 3) the proportion of an ecoregion’s remaining habitat contained within the site; and 4) level of threat. We evaluate these elements in three ways: individually; combined into an importance ranking (including elements 1-3); and an overall threat-weighted ranking (combining all four elements). We illustrate the flexibility of this approach by demonstrating how results can be filtered by ecoregion, country, biome, realm, or globally. We also examine how appropriate recognition of Indigenous and Traditional Territories (ITTs) could secure the protection of many Conservation Imperatives Sites.

This prioritization addresses a growing concern among conservation biologists: that increased efforts to meet the 2030 deadline for protecting 30% of land and sea could lead to a focus on the quantity of land protected—in order to reach a numerical target—rather than quality, the latter achieved by first selecting the most ecologically important sites (Geldmann, 2026). Current coverage of protected and conserved areas—the term used for the combination of protected areas and other effective area-based conservation measures (OECMs)—is 18.43% for terrestrial and inland waters as of April 2026 (Protected Planet, 2026). Therefore, the fast-approaching deadline to meet the 30% coverage target by 2030 makes protecting the most sensitive areas a matter of urgency. Parties to the CBD have worked to define national targets as contributions to the global target, and many are planning to expand their systems of protected areas and OECMs. As a result, this prioritization can provide much-needed science-based guidance.

## Materials and methods

### Source Data for Conservation Imperatives Sites

Conservation Imperatives Sites were identified by Dinerstein et al. (2024). Located primarily in tropical and subtropical moist forest biomes, they are sites that support rare, narrow-range endemic, and endangered species. The 16,825 Conservation Imperatives Sites originally identified cover 1.22% of Earth’s land surface. These sites were delineated by combining six widely used data layers, including the Alliance for Zero Extinction (AZE) sites, Key Biodiversity Areas (KBAs) and derivatives (a range-size rarity raster and a small-ranges vertebrates raster) from the IUCN Red List species range data (Dinerstein et al., 2024). They erased protected areas from this spatial layer, ensuring all Conservation Imperative Sites fall outside formal protected area networks (see Dinerstein et al. (2024) for further details).

We updated the Conservation Imperatives layer to remove any protected and conserved areas added to the World Database on Protected Areas (WDPA) and World Database on OECMs (WD-OECM) (UNEP-WCMC and IUCN, 2025) since the 2024 publication. In this step, we excluded proposed protected areas and OECMs, as well as sites where the designation status is unreported. We also excluded UNESCO Man and Biosphere Reserves (MAB) sites as these often include buffer and transition zones lacking protected area status (Protected Planet, 2024). This refinement resulted in a total of 16,646 Conservation Imperatives Sites that served as the basis for the prioritization.

We used the public version of the WDPA, which omits restricted data. Restricted data accounts for the majority of protected areas in China, India, Türkiye and Saint Helena, with smaller numbers restricted in Estonia, Lebanon, Ireland, and New Zealand. As a result, some Conservation Imperatives Sites in these countries may already fall within protected areas.

### Data used to prioritize Conservation Imperatives Sites

Besides data on Conservation Imperatives Sites and protected and conserved areas, four other datasets were incorporated into the prioritization analysis (Table 1).

**Table 1:**
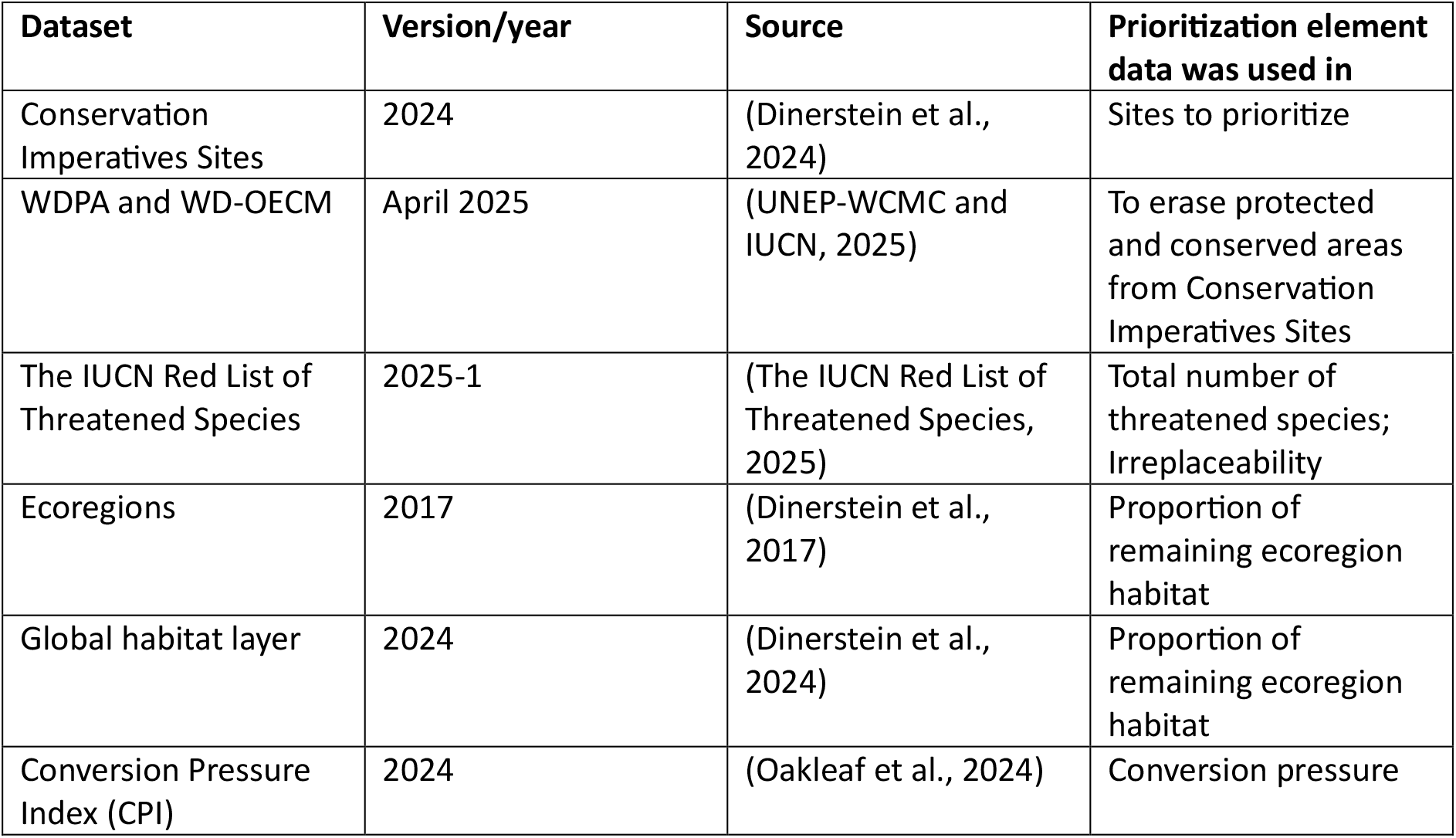
Datasets used in the prioritization analysis.

### Prioritization approach

Our framework combines four important elements to prioritize Conservation Imperatives Sites for protection. These are: (1) the total number of threatened species (critically endangered, endangered and vulnerable) present at the site; (2) irreplaceability of the site; (3) the proportion of an ecoregion’s remaining habitat represented by the site, emphasizing sites that represent the last opportunities to conserve remaining habitat supporting geographically distinct assemblages of biodiversity; and (4) the level of conversion pressure at the site.

#### 1. Threatened Species

The ultimate goal is to prevent extinctions; therefore, sites with the greatest number of threatened species are prioritized. We use species ranges from the IUCN Red List (The IUCN Red List of Threatened Species, 2025) for terrestrial vertebrates (mammals, birds, reptiles and amphibians). These taxa have been comprehensively assessed which avoids geographical biases and provides a good proxy for overall biodiversity. Only extant species (presence = 1) were included. Species ranges were intersected with the Conservation Imperatives Sites, and the number of unique species were summed by site, ecoregion, country, biome and realm.

#### 2. Irreplaceability

In addition to the total number of threatened species, the degree to which species are dependent on a given site is also a critical differentiating factor. For this assessment, we applied the irreplaceability measure described in Le Saout et al. (2013). This approach calculates the percentage of each species’ range that overlaps Conservation Imperatives Sites, with sites being more irreplaceable if they have more species with a higher dependence on them (as measured by the percentage of their range within the site). The irreplaceability score thus reflects both the degree of species dependence on the site, and the total number of species found there (see Le Saout et al. (2013) for further details). We use species ranges of mammals, birds, amphibians and reptiles from the 2025-1 version of The IUCN Red List of Threatened Species data; this step therefore serves as an irreplaceability analysis for terrestrial vertebrates.

#### 3. Remaining Habitat

In heavily converted ecoregions, particularly grassland ecoregions and those representing other non-forest biomes, little natural habitat remains inside or outside protected and conserved areas. Conservation Imperatives Sites that represent some of the last opportunities to conserve the characteristic habitats, vegetation, species assemblages, and species interactions within these ecoregions are therefore also prioritized. To quantify remaining habitat within each ecoregion, we intersected ecoregions data (Dinerstein et al., 2017) with a global habitat map developed following the methodology of Dinerstein et al. (2024). This layer was then intersected with the Conservation Imperatives Sites and the proportion of remaining ecoregion habitat within each site was calculated.

#### 4. Conversion Pressure

Assuming feasibility of protection is not an overriding constraint, sites facing higher levels of threat require protection sooner than less disturbed areas. We use predicted conversion pressure by 2030, rather than current threat levels, because protection is not immediate and we sought to prioritize sites likely to experience high pressure in the near future. We treat conversion pressure as a separate variable from the IUCN threat category as threats to an area can accelerate within months (e.g., exploration or discovery of oil, gas, or mineral deposits), rapidly increasing pressure on a site. To assess predicted future pressure on Conservation Imperatives Sites, we use the Conversion Pressure Index (CPI) which combines projected increases in human modification to 2030 with industrial development suitability (Oakleaf et al., 2024). We calculate the median CPI score for each Conservation Imperative Site and classify sites into six threat categories following those defined in Oakleaf et al. (2024): very low (CPI ≤ 0.059); low (0.059 < CPI ≤ 0.265); moderate low (0.265 < CPI ≤ 0.471); moderate high (0.471 < CPI ≤ 0.677); high (0.677 < CPI ≤ 0.883); very high (0.883 < CPI ≤ 1.000).

Eighteen Conservation Imperatives Sites did not overlap with a CPI grid cell. These were small coastal sites falling just beyond the CPI layer extent and were assigned the average CPI score of other sites in their ecoregion. One site was the only site in its ecoregion so was assigned the average of the sites within its biome.

### Ranking sites

To derive an overall ranking of sites — which can be used at any level (ecoregion, country, biome, realm, global) — we first combine variables 1-3 (site importance) and then weight this by variable 4 (threat level). We do this because: 1) variables 1-3 are unlikely to change significantly over the next 5-10 years at most sites, whereas threat level could change rapidly; and 2) it gives users the flexibility to apply no threat weighting or a modified one.

To combine the importance variables, we first ranked sites separately for each variable; these ranks were then summed to produce an overall importance ranking. To make rankings easier to compare, we scaled them to a 0–1 range.

To account for the level of threat at each site, we multiplied the scaled overall importance ranking by a weight based on the threat categories. The threat weight was calculated using the formula:

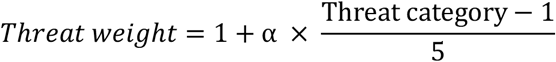

where α is the maximum proportional increase in threat weighting. We use *α* = 1, meaning the highest weight is double the lowest. Using this formula, the threat categories translate into the following weightings: very low = 1; low = 1.2; moderate low = 1.4; moderate high = 1.6; high = 1.8; and very high = 2. The α value can be changed to adapt the weightings.

We then group the sites into 10 quantiles based on the threat-weighted rankings, with the first quantile group (highest-priority group) comprising sites in most urgent need of protection. The rationale here is that 1) there will be little difference between closely ranked sites, and 2) it gives nations, funders and NGOs defined goals to target for immediate action.

### Indigenous and traditional territories

Although KMGBF Target 3 includes the wording ‘Indigenous and traditional territories’, there is currently no agreed definition for these territories in that context. For this analysis, we interpret them as encompassing territories and areas that are owned, governed, or used by Indigenous Peoples and local communities. To produce a spatial layer of Indigenous and traditional territories, we merged two datasets from LandMark: Indicative Areas of Indigenous and Community Land Rights (LandMark, 2024a) and Indigenous Peoples’ and Local Community Lands and Territories (LandMark, 2024b).

We then intersected this layer with the Conservation Imperatives Sites to identify the number of Conservation Imperatives Sites that overlap at least partially with Indigenous and traditional territories. Further analysis of these overlaps was not undertaken because the Indigenous and traditional territories dataset is not yet globally representative.

## Results

### Threatened Species

We found that 97.54% of Conservation Imperatives Sites (16,236 sites) contain at least one threatened species of mammal, bird, reptile or amphibian. In addition, 88.50% of Conservation Imperatives Sites (14,731 sites) contain at least ten threatened species. Refining further, 1.97% (324 sites) contain >50 threatened species threatened species of mammal, bird, reptile or amphibian. The median number of threatened species per site is 21.

### Irreplaceability

Applying the irreplaceability analysis finds that 33.46% of Conservation Imperatives Sites (5,569 sites) are more irreplaceable for terrestrial vertebrates than 90% of protected and conserved areas in the WDPA and WD-OECM (298,042 sites); 14.63% of Conservation Imperatives Sites (2,436 sites) are more irreplaceable than 99% of protected and conserved areas. The median irreplaceability value for sites is 0.00005.

### Remaining Habitat

The proportion of remaining ecoregion habitat found within Conservation Imperatives Sites varies widely. One site in the Admiralty Islands lowland rain forests ecoregion (Australasia) contains 90.13% of the remaining habitat of that ecoregion. However, the median value across all sites is 0, owing to the small size of many Conservation Imperatives Sites and the correspondingly small proportion of remaining ecoregion habitat they contain.

### Conversion Pressure

We found that 48.59% of Conservation Imperative Sites are predicted to have high or very high conversion pressure by 2030; only 3.68% of sites are predicted to experience low or very low conversion pressure. A further 10.74% and 36.99% are predicted to experience moderate low and moderate high conversion pressure, respectively. The median CPI value of sites is 0.671 (moderate high).

### Overall rankings

We prioritize 1,667 sites — sites in the highest-priority quantile group — that are in most urgent need of protection (see Supplementary Material 1 for the full list of sites). These 1,667 sites span 501,426 km^2^, covering 0.37% of Earth’s land surface. The median size of sites in the highest-priority group is 40.78 km^2^ (range: 0.12 - 37,173 km^2^), with 31.91% being over 100 km^2^. Most sites are tropical, as well as some notable clusters around the Mediterranean and in southern Africa (Figure 1). The highest-priority sites are found in 71 countries. However, 71.45% of the sites are restricted to 10 countries and 87.34% of the sites occur in only 20 countries. The Philippines, Indonesia, India, and Madagascar alone contain 49.55% of the sites in the highest-priority group. Some sites in this group are large and mostly connected, such as sites in the Sulawesi lowland and montane rain forest ecoregions of Indonesia. Other sites are smaller and fragmented such as those in the Eastern Arc forests and Northern Swahili coastal forest ecoregions of Tanzania, and the Bahia interior forests ecoregion of eastern Brazil (Figure 1). Of the 1,667 highest priority sites, 51.53% (859 sites) are within 2.5 km^2^ of an existing protected area.

**Figure 1.**
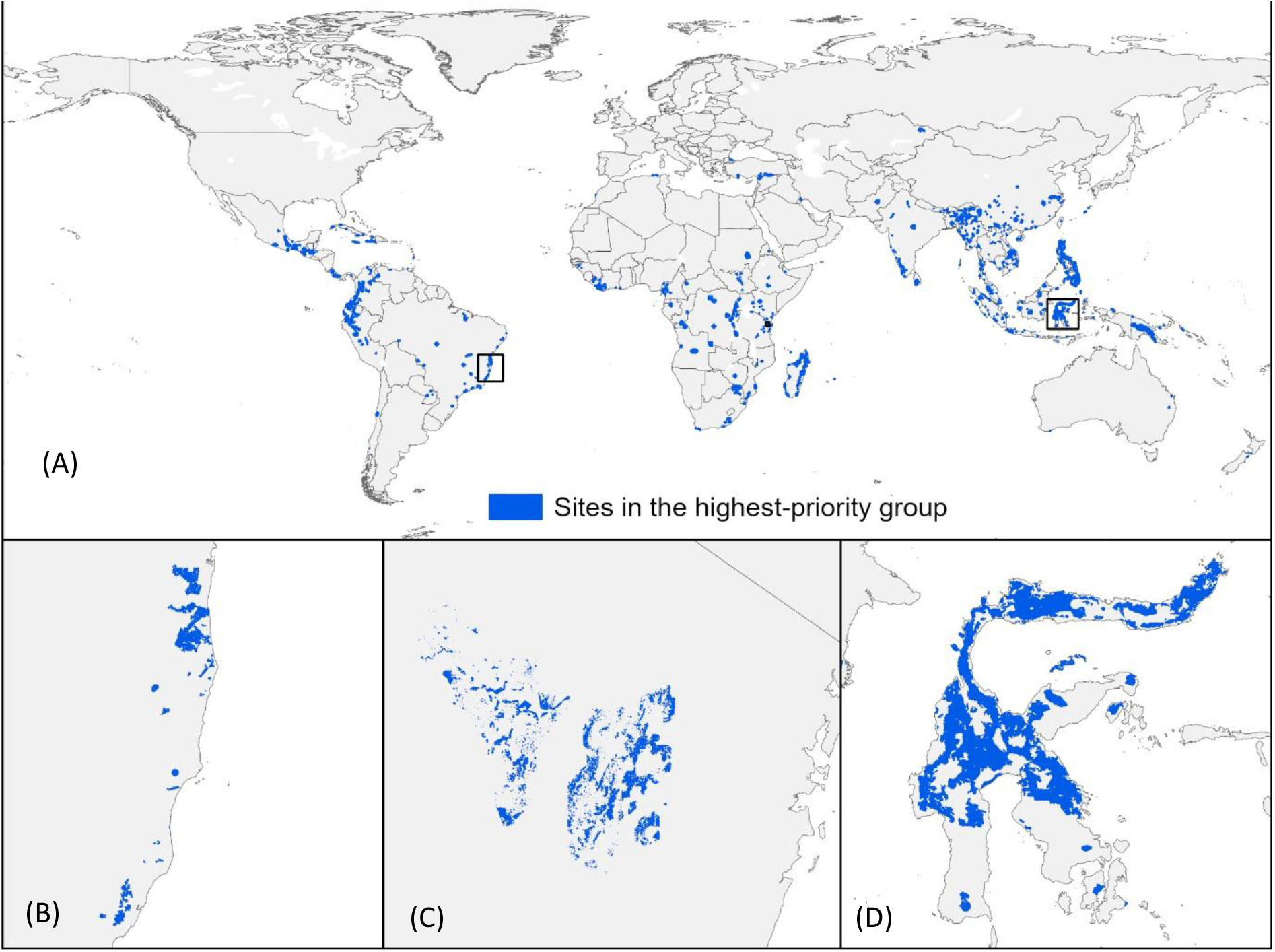
(A) Global distribution of the 1,667 Conservation Imperatives Sites in the highest-priority group (sites have been made to look larger than they are to be more visible on the global map). Insets for (B) the Bahia interior forests of eastern Brazil; (C) the Eastern Arc forests and Northern Swahili coastal forests of Tanzania; and (D) the Sulawesi lowland and montane rain forests of Indonesia portray the actual spatial extent of the Conservation Imperatives Sites.

Individual variable rankings are also broken down into quantile groups to examine variation among countries. Madagascar contains the most sites (306) in the top quantile group for threatened species, with Indonesia having the most sites in the top quantile group for irreplaceability (166 sites), remaining ecoregion habitat (135 sites), and conversion pressure (669 sites) (Figure 2 and Table 2). However, the Philippines has the greatest number of sites in the overall highest-priority group (368), followed by Indonesia (206), India (149), Madagascar (103) and Brazil (75) (Table 2). These five countries also have the greatest number of Conservation Imperatives Sites in total.

**Table 2.** Top 10 countries with the most Conservation Imperatives Sites in the highest-priority group. The number of Conservation Imperatives Sites in the top quantile group of rankings for each individual variable and combined importance rank are also shown. See Supplementary Material 2 for the full list of countries.

| Country | Total number of Conservation Imperatives Sites | Number of sites in top group for threatened species | Number of sites in top group for irreplaceability | Number of sites in top group for ecoregion habitat | Number of sites in top group for CPI | Number of sites in top group for combined importance rank | Number of sites in highest-priority group for overall threat weighted rank |
| --- | --- | --- | --- | --- | --- | --- | --- |
| Philippines | 3418 | 274 | 140 | 130 | 513 | 324 | 368 |
| Indonesia | 1960 | 212 | 166 | 135 | 669 | 162 | 206 |
| India | 371 | 204 | 92 | 123 | 0 | 188 | 149 |
| Madagascar | 941 | 306 | 66 | 24 | 24 | 104 | 103 |
| Brazil | 3286 | 14 | 81 | 40 | 175 | 55 | 75 |
| Colombia | 687 | 42 | 100 | 59 | 8 | 68 | 64 |
| Ecuador | 626 | 93 | 54 | 31 | 66 | 54 | 61 |
| Myanmar | 102 | 91 | 21 | 31 | 13 | 65 | 59 |
| China | 256 | 67 | 38 | 43 | 3 | 67 | 53 |
| Mexico | 225 | 32 | 78 | 81 | 1 | 47 | 48 |

**Figure 2.**
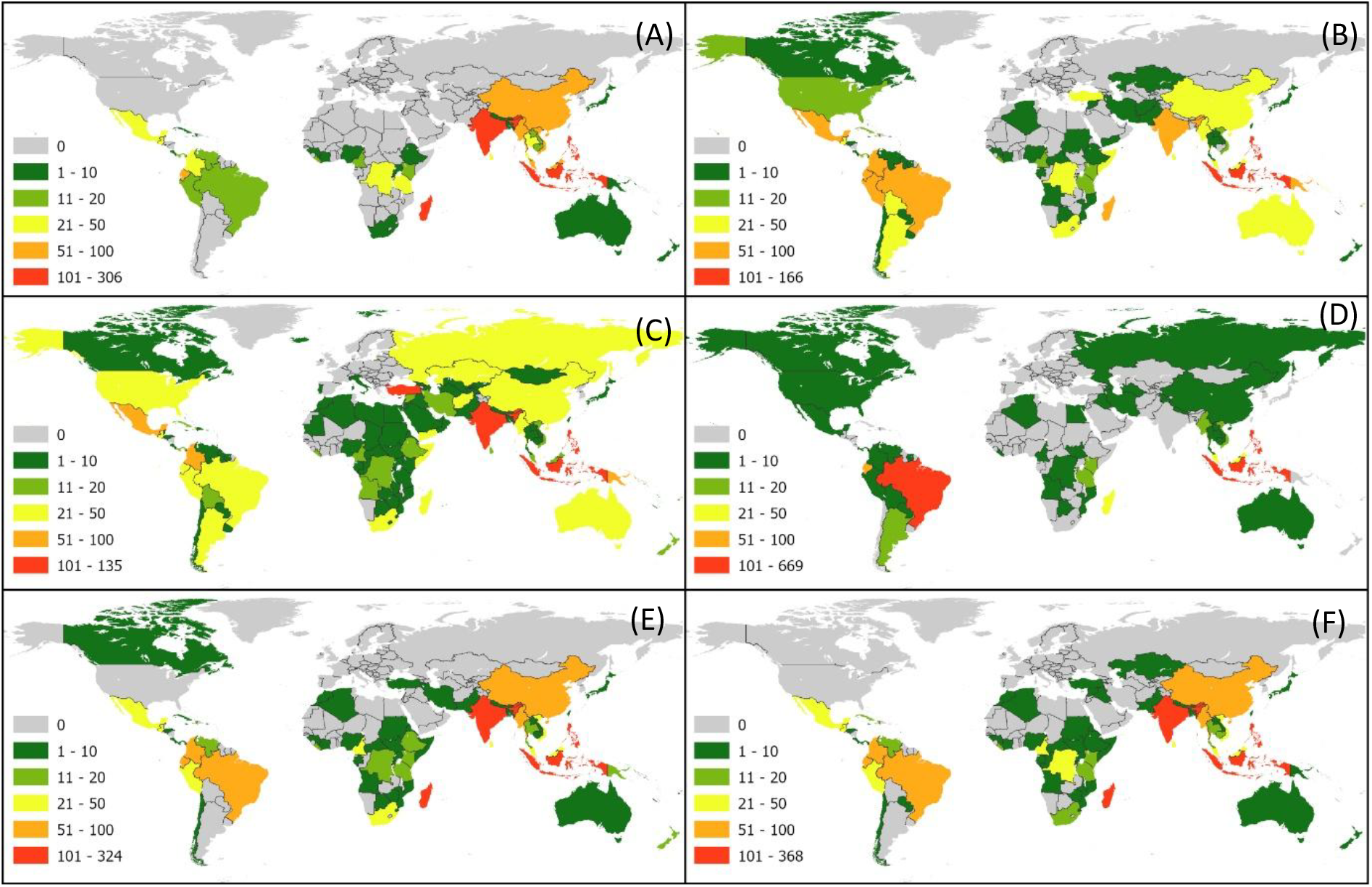
Number of Conservation Imperatives Sites in the top quantile group of rankings for (A) number of threatened species; (B) irreplaceability; (C) percent remaining ecoregion habitat; (D) conversion pressure index; (E) combined importance ranking; and (F) overall threat weighted ranking (highest-priority sites) by country.

Brunei Darussalam has the highest median number of threatened species (64) in Conservation Imperatives Sites; however, the country has only one Conservation Imperatives Site. Malaysia has the second highest median number of threatened species (58) and has 73 Conservation Imperatives Sites in total. Countries with high median irreplaceability scores generally only have a few Conservation Imperatives Sites in total, with Mayotte (one site), Dominica (two sites) and Comoros (three sites) ranking as the top three. Similarly, the same three countries occupy the top three positions for remaining ecoregion habitat. Conservation Imperatives Sites in Lao People’s Democratic Republic have the highest median Conversion Pressure Index score followed by Malaysia and Brunei Darussalam. Liberia has the highest median overall threat-weighted ranking, with the country’s 22 sites having a median ranking of 514.5, followed by Guatemala and Sri Lanka (Table 3)

**Table 3.** Top 10 countries with the highest median threat weighted rank. The median values for each individual variable are also shown. See Supplementary Material 2 for the full list of countries.

| Country | Total number of Conservation Imperatives Sites | Median number of threatened species | Median irreplaceability score | Median percent of ecoregion habitat | Median CPI score | Median importance rank | Median threat weighted rank |
| --- | --- | --- | --- | --- | --- | --- | --- |
| Liberia | 22 | 33.5 | 0.0215 | 0.265 | 0.767 | 590.5 | 514.5 |
| Guatemala | 40 | 32 | 0.0325 | 0.285 | 0.714 | 928.5 | 575.5 |
| Sri Lanka | 49 | 47 | 0.0121 | 0.09 | 0.633 | 608 | 841 |
| Cambodia | 19 | 42 | 0.0001 | 0.03 | 0.806 | 1981 | 1033 |
| Mayotte | 1 | 17 | 2.5545 | 16.75 | 0.747 | 1530 | 1064 |
| Lao People's Democratic Republic | 19 | 41 | 0.0002 | 0.02 | 0.877 | 2458 | 1095 |
| Guinea-Bissau | 1 | 23 | 0.0037 | 0.07 | 0.736 | 1835 | 1304 |
| Saint Lucia | 3 | 15 | 0.1918 | 8.99 | 0.714 | 1943 | 1387 |
| Myanmar | 102 | 39 | 0.0006 | 0.06 | 0.638 | 1055 | 1406 |
| Malaysia | 73 | 58 | 0.0046 | 0.01 | 0.854 | 1056 | 1504 |

Australasia contains the highest number of sites (592 sites) in the highest-priority group, with 90% (533 sites) of these in three ecoregions (Sulawesi lowland rain forests, Central Range Papuan montane rain forests and Sulawesi montane rain forests). The Neotropics follows closely with 557 sites, however these are distributed among a greater number of ecoregions with 16 ecoregions having more than 10 sites in the highest-priority group (compared to three ecoregions in Australasia) (Table 4).

**Table 4.**
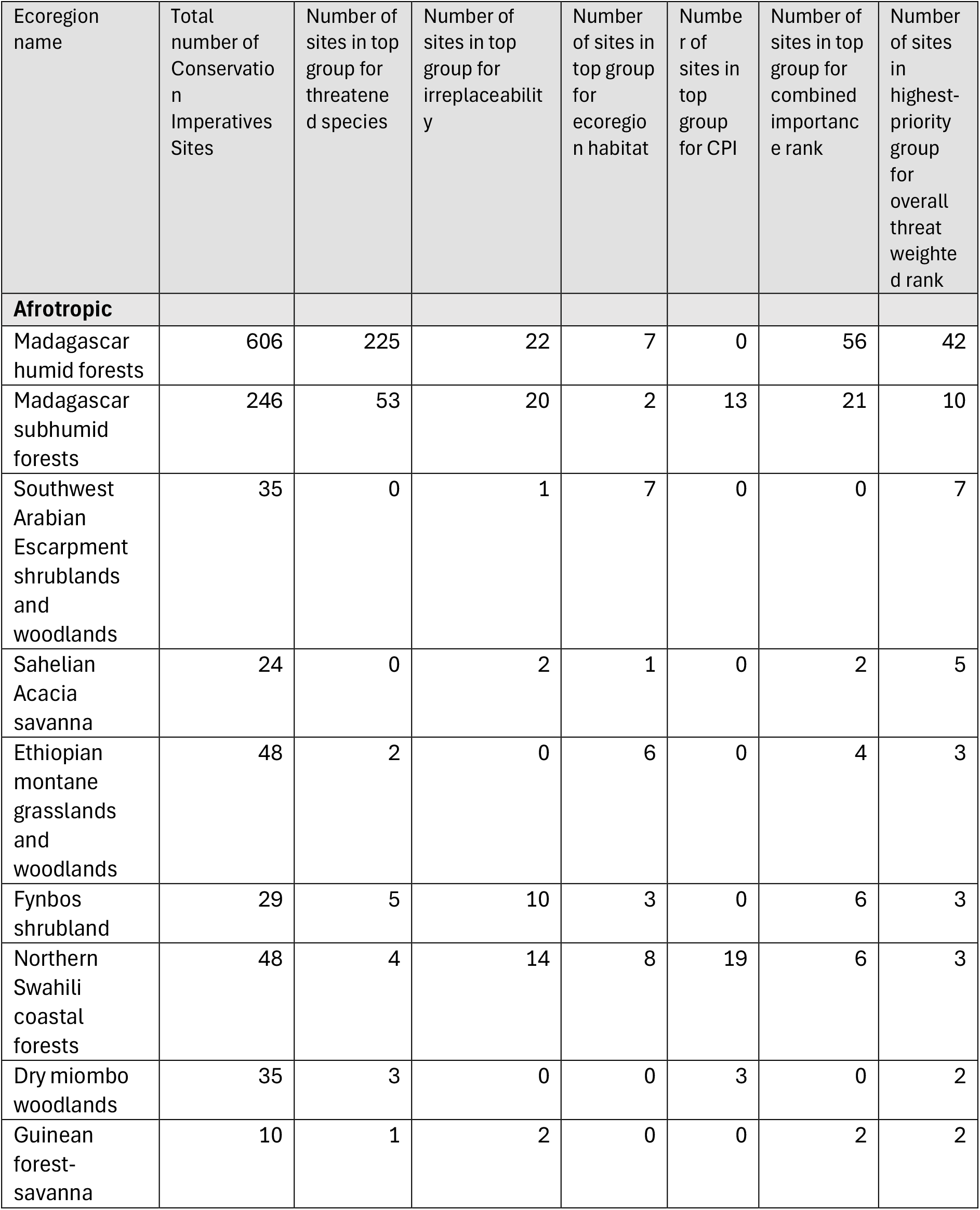
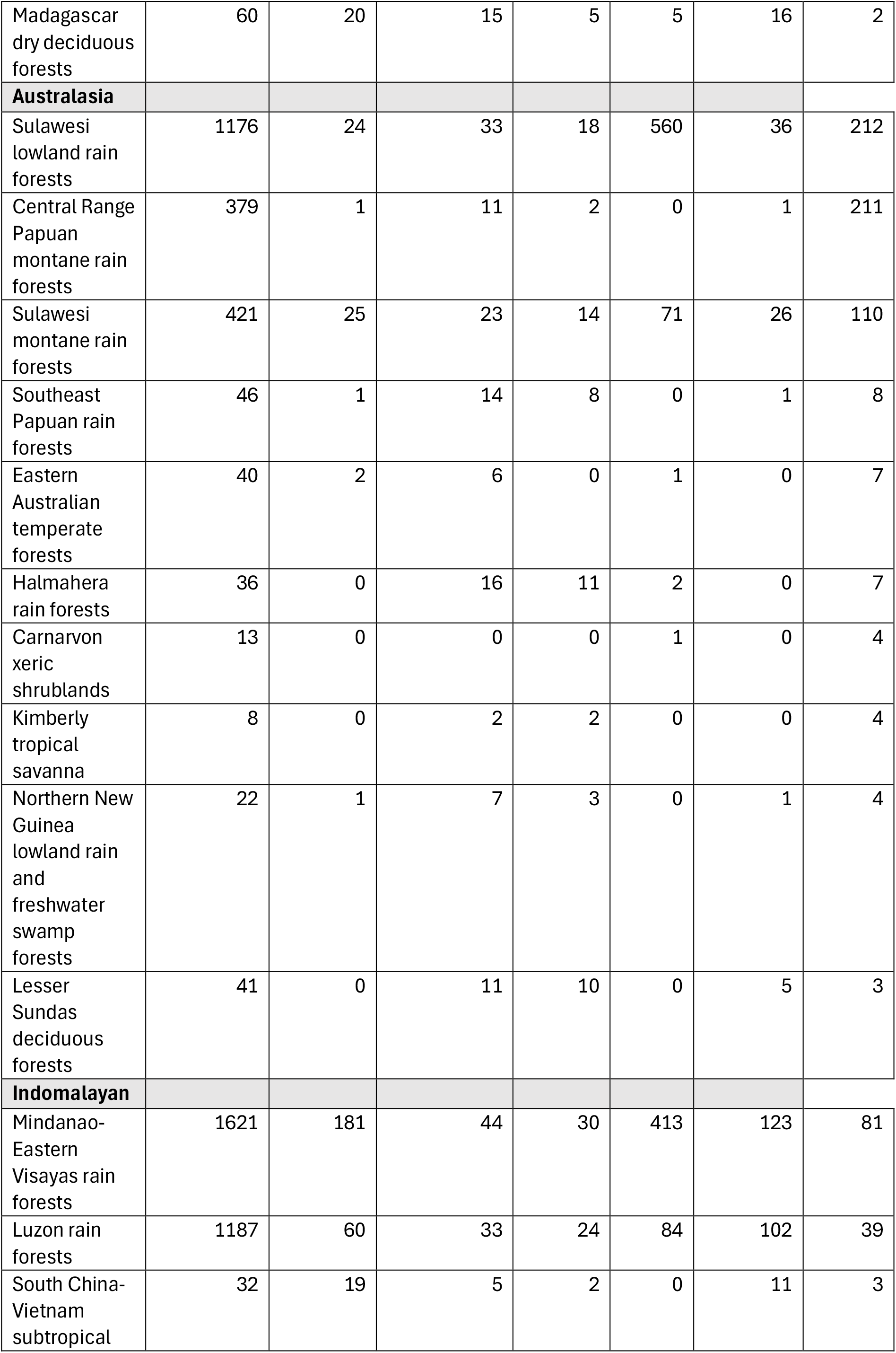
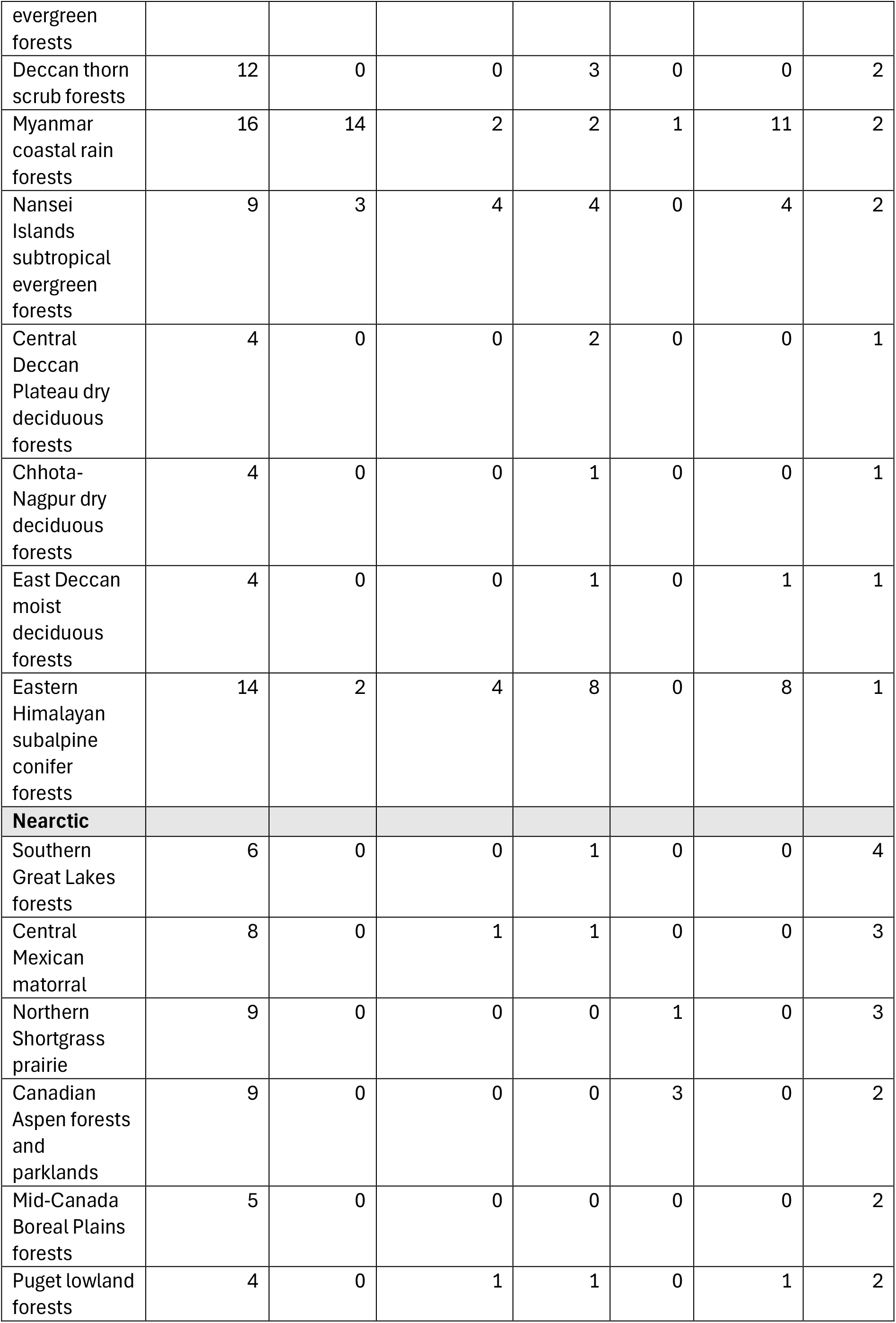
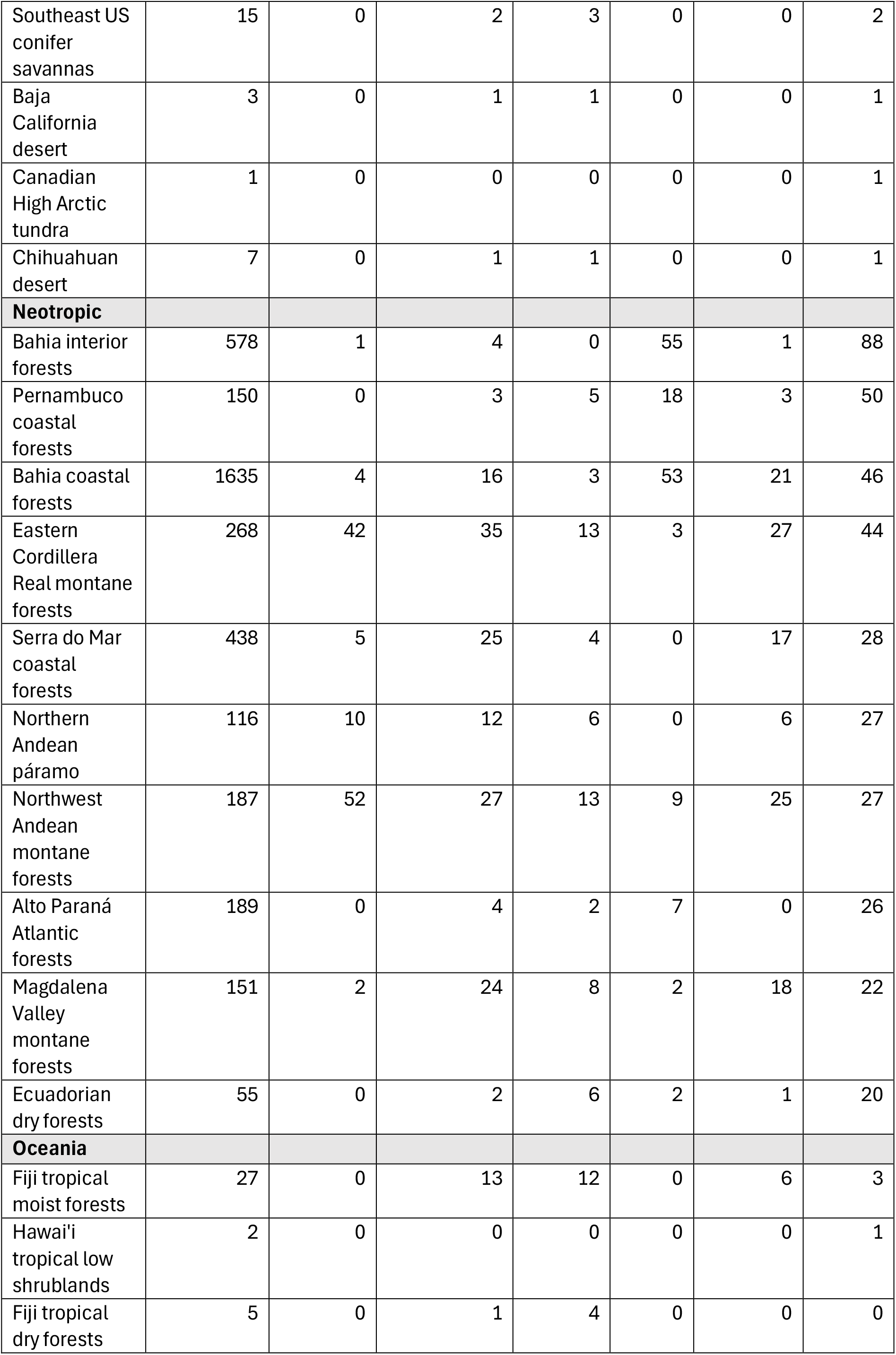
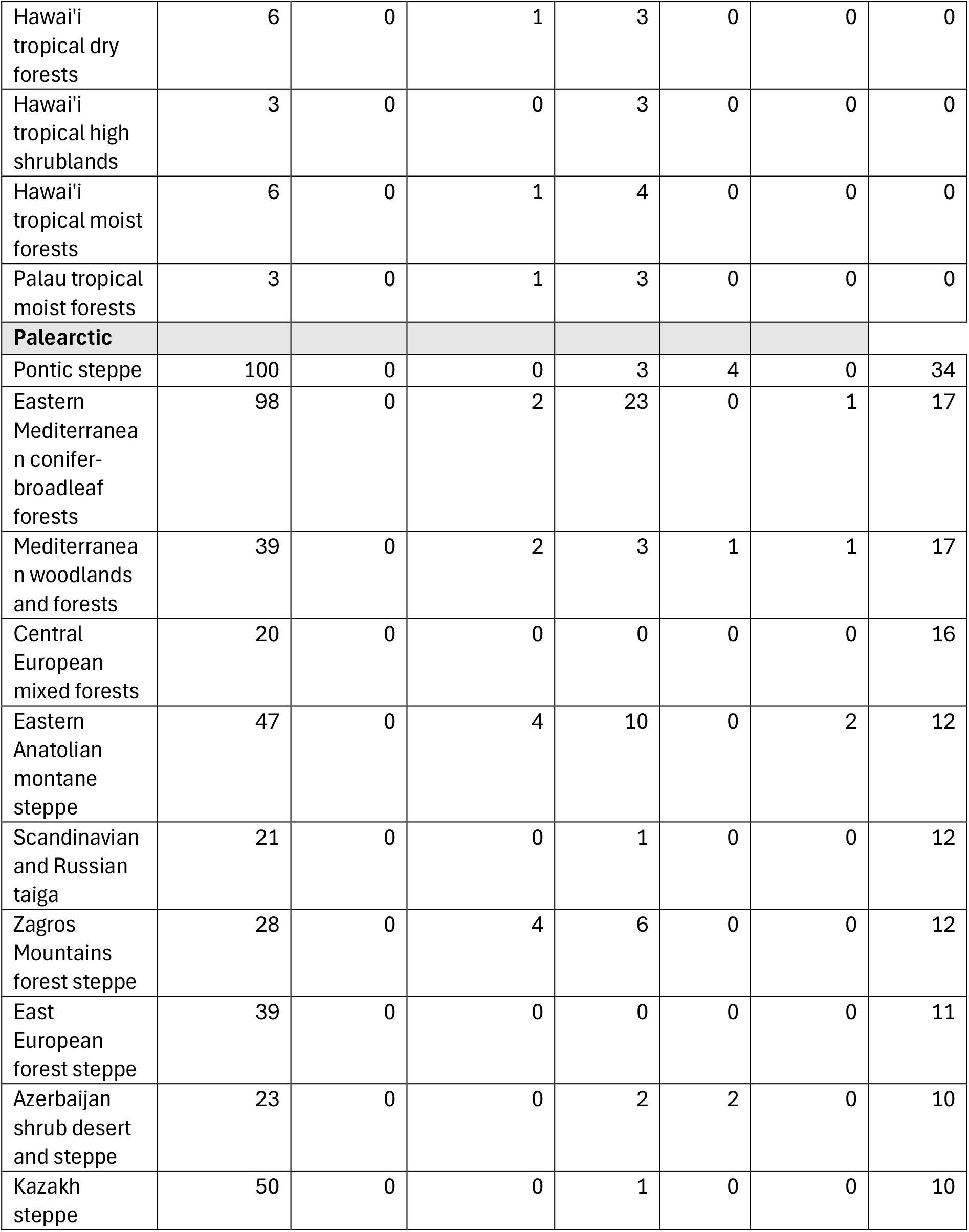
Top 10 ecoregions in each realm with the most Conservation Imperatives Sites in the highest-priority group. Number of Conservation Imperatives Sites in the top quantile group for each of the variables is also shown. Oceania only has seven ecoregions with sites in the top quantile group of any of the variables, thus only seven are displayed (see Supplementary Material 3 for the full list of ecoregions, biomes and realms).

**Table 5.** The top 10 ranked Conservation Imperatives Sites in Guatemala and the individual variable scores.

| National threat weighted rank | Global threat weighted rank | Ecoregion | Size (km<sup>2</sup>) | Number of threatened species | Irreplaceability score | Percent of ecoregion habitat | CPI category |
| --- | --- | --- | --- | --- | --- | --- | --- |
| 1 | 40 | Central American pine-oak forests | 1052.94 | 53 | 7.41 | 3.88 | High |
| 2 | 83 | Central American montane forests | 64.98 | 42 | 0.27 | 1.12 | High |
| 3 | 85 | Chiapas montane forests | 50.33 | 50 | 0.04 | 1.95 | High |
| 4 | 93 | Petén-Veracruz moist forests | 195.05 | 52 | 1.53 | 0.28 | High |
| 5 | 101 | Motagua Valley thorn scrub | 61.60 | 42 | 0.02 | 63.68 | High |
| 6 | 111 | Central American pine-oak forests | 104.85 | 42 | 0.30 | 0.39 | High |
| 7 | 114 | Central American pine-oak forests | 110.69 | 55 | 0.06 | 0.41 | High |
| 8 | 118 | Sierra Madre de Chiapas moist forests | 194.21 | 32 | 0.11 | 3.95 | High |
| 9 | 154 | Central American Atlantic moist forests | 121.76 | 38 | 0.05 | 0.35 | High |
| 10 | 159 | Central American Atlantic moist forests | 455.57 | 29 | 1.31 | 1.29 | High |

### National Level results - Guatemala

Guatemala offers an excellent example of how our prioritization results can be applied at a national level. We choose Guatemala because, although it contains only 40 Conservation Imperatives Sites in total, 29 are in the highest-priority group of sites in most urgent need of protection and have a high median threat-weighted rank. In addition, 29 sites have high predicted future conversion pressure, while the remaining 11 have moderately high conversion pressure.

The highest-ranked site in Guatemala (ranked 40th globally) is a large site (1,052.94 km^2^) in the Central American pine-oak forests ecoregion. It is home to 53 threatened terrestrial vertebrates, has a very high irreplaceability score, and retains 3.88% of the remaining habitat of its ecoregion. It also has high predicted future conversion pressure (Table 3). Most sites in Guatemala score highly overall, but examining individual variable scores reveals the site ranked fifth highest nationally as a potential priority. It holds 63.68% of the remaining habitat of the Motagua Valley thorn-scrub ecoregion and is experiencing high conversion pressure, factors that may justify its prioritization for national protection.

### Indigenous and Traditional Territories

At least 13.37% (2,225 sites) of Conservation Imperative Sites are partially or fully within Indigenous and traditional territories. Of the sites that are in the highest-priority group, at least 15.24% (254 sites) are partially or fully within Indigenous and traditional territories.

## Discussion

### Prioritizing the Highest Quality Unprotected Locations of Rarity

Although protecting all Conservation Imperatives Sites is a global priority, limited conservation funding and increasing pressures on biodiversity make prioritization essential. Our assessment elevates 1,667 of the total 16,825 sites for highest consideration based on biodiversity value and threat. Several key findings form the basis for an emerging strategy to efficiently protect rare nature. First, more than one-third of Conservation Imperatives Sites have higher irreplaceability than 90% of current protected and conserved areas. Second, almost half of Conservation Imperatives Sites are predicted to experience high or very high conversion pressure by 2030, emphasizing the urgency of targeted protection. Third, the 1,667 highest priority sites are concentrated geographically: 71% occur in only 10 countries and 87% are distributed among 20 countries. The Philippines, Indonesia, India, and Madagascar combined contain almost half of all highest priority sites. Finally, these 1,667 sites comprise just 0.37% of the Earth’s terrestrial surface, highlighting a relatively small area requiring immediate action. These findings suggest that focusing effort and establishing new protected and conserved areas in around 20 countries over a relatively limited land area could provide an efficient next step in implementing the KMGBF’s Target 3, combatting extinctions and loss of unique habitats.

Under the KMGBF, Target 3 emphasizes increasing coverage to 30% but places equal emphasis on ensuring the quality of protected and conserved areas (CBD, 2022). Careful prioritizations such as ours ensure that protected and conserved areas are expanded to the highest-quality areas. Our prioritization framework enables planners supporting governments, NGOs and philanthropic funders to identify the most important and threatened sites. Finally, the framework offers flexibility to prioritize by importance variables (Threatened Species, Irreplaceability, Remaining Habitat) individually, combined or weighted by threat (Conversion Pressure), making it easily adapted to local, national, or regional contexts.

### Mechanisms for Conserving Rarity and Preventing Extinctions

Through the KMGBF, Parties to the CBD and other actors recognise the importance of protected areas and OECMs for halting biodiversity loss. Evidence shows that protected areas can prevent biodiversity loss when effectively managed (Cazalis et al., 2020; Wauchope et al., 2022; Cooke et al., 2023). OECMs provide additional pathways for recognizing governance systems that already deliver conservation benefits, potentially empowering local communities and improving access to funding (Alves-Pinto et al., 2021). Although studies on OECM effectiveness remain limited (Cook, 2023), OECMs are expected to contribute substantially toward meeting the KMGBF’s targets. Equitable and effective governance of protected and conserved areas is also increasingly recognized as fundamental (e.g. Eklund & Cabeza, 2017; Huber et al., 2023; Mast et al., 2025).

Indigenous Peoples and local communities play a pivotal role in sustaining biodiversity (Dinerstein et al., 2020; WWF et al., 2021). As highlighted in the Protected Planet Report 2024 (UNEP-WCMC and IUCN, 2024), appropriate recognition of Indigenous and traditional territories (ITTs) as protected areas, OECMs, or through additional pathways under discussion within CBD processes could greatly advance Target 3 implementation. We find that a substantial proportion (at least 13%) of Conservation Imperatives Sites overlap with ITTs. Although these proportions may appear modest, it likely underestimates the true overlap due to incomplete mapping of ITTs. Appropriately recognizing these areas could secure the rights of Indigenous Peoples and local communities and deter external pressures.

Designating protected areas and recognising existing governance systems that conserve Conservation Imperatives Sites have a clear role to play. However, new protected and conserved areas alone, even when effectively managed, will not prevent some threatened species from going extinct. Threats such as invasive species, disease, and climate change often require interventions beyond area-based protection. Invasive species are major drivers of extinctions on islands, necessitating eradication and biosecurity programmes (Hoffmann et al., 2010; Spatz et al., 2017; Dinerstein et al., 2024). Diseases such as chytridiomycosis threaten amphibians globally, with restrictions on long-distance trade currently the best defence (Kriger and Hero, 2009; Garner et al., 2016). Climate change will increasingly drive extinctions unless emissions are reduced and land management enables species range shifts (Warren et al., 2013; Urban, 2015; Román-Palacios and Wiens, 2020; Parks et al., 2023).

Small, protected sites are often insufficient to prevent species loss, as many species require larger, connected habitats (Pimm and Jenkins, 2019). In such cases, it may be necessary to undertake restoration, expand protected and conserved areas, or where relevant, integrate Indigenous and traditional territories into broader conservation networks, while respecting rights. However, some species—particularly small animals and some plants in ancient, highly endemic systems or on islands—can persist long-term in small habitat patches if ecological conditions are maintained (e.g. millepedes, spiders and African violets in the Eastern Arc Mountains of Tanzania (Dimitrov et al., 2012; Enghoff et al., 2025; Malumbres-Olarte et al., 2025). If these ecological conditions are disrupted though, it can lead to rapid population declines and extinctions (e.g. the Kihansi spray toad (Channing et al., 2006)). These findings highlight the importance of protecting small sites with small ranged endemic species as well protecting and connecting larger landscapes.

As with all global analyses, validation by national experts is necessary to ensure appropriate local prioritization. Our framework allows for the substitution of global data with national data where available. For example, data on the distribution and status of taxa lacking comprehensive assessments by the IUCN Red List (such as many vascular plants) could be incorporated into the prioritization where such national scale data exists. The threat weighting can also be easily adjusted to local or regional scales (code is available at https://doi.org/10.5281/zenodo.19368637). Cost and feasibility are critical variables that affect prioritization but are best incorporated at the local scale rather than inferred from global datasets. These factors should be considered alongside biological importance and threat. Proximity to existing protected and conserved areas may also be relevant, especially with over half of the highest priority sites being within 2.5 km^2^ of an existing protected area, as expansion may be more feasible or cost-effective than establishing new reserves (Dinerstein et al., 2024). Restoring habitat between Conservation Imperatives Sites and nearby protected and conserved areas could also be an effective and strategic use of investments in restoration.

### Conclusion

Nations should undertake national-level prioritizations to ensure protection of the most important biodiversity areas, while incorporating globally significant sites such as Conservation Imperatives in order to meet the KMGBF targets (Plumptre et al., 2024). Our prioritization framework can be adapted to local, national, or regional contexts and can help governments, funders and NGOs focus efforts on the most biologically important and threatened Conservation Imperatives Sites. Of the 16,825 Conservation Imperatives Sites, we elevate 1,667 that require the most urgent protection. Many Conservation Imperatives Sites are more irreplaceable than existing protected and conserved areas, and nearly half are projected to face high conversion pressure by the end of the decade. Targeting conservation actions in a relatively small number of countries could deliver major conservation benefits over the next five years. Protecting these sites is critical for preventing extinctions and they are strong candidates for protected and conserved area expansion under Target 3, adding quality to quantity on the path to reach 30%, alongside recognition of Indigenous Peoples’ and local communities’ land rights.

## Acknowledgments

We thank S. Pimm for helpful comments on the draft manuscript.

## Notes

### Competing Interest Statement

This study received funding from the non‐profit organization Art into Acres (https://www.arttoacres.org), which HM is affiliated with.

### Funding

- Art into Acres
- Nature Data Lab

